# Ultrafast clustering of single-cell ow cytometry data using FlowGrid

**DOI:** 10.1101/394189

**Authors:** Xiaoxin Ye, Joshua W. K. Ho

**Affiliations:** Victor Chang Cardiac Research Institute, Sydney, Australia; University of New South Wales, Sydney, Australia; School of Biomedical Sciences, Li Ka Shing Faculty of Medicine, The University of Hong Kong, Hong Kong, China

**Keywords:** Clustering, flow cytometry, single cell, DBSCAN

## Abstract

Flow cytometry is a popular technology for quantitative single-cell profiling of cell surface markers. It enables expression measurement of tens of cell surface protein markers in millions of single cells. It is a powerful tool for discovering cell sub-populations and quantifying cell population heterogeneity. Traditionally, scientists use manual gating to identify cell types, but the process is subjective and is not effective for large multidimensional data. Many clustering algorithms have been developed to analyse these data but most of them are not scalable to very large data sets with more than ten million cells.

Here, we present a new clustering algorithm that combines the advantages of density-based clustering algorithm DBSCAN with the scalability of grid-based clustering. This new clustering algorithm is implemented in python as an open source package, FlowGrid. FlowGrid is memory efficient and scales linearly with respect to the number of cells. We have evaluated the performance of FlowGrid against other state-of-the-art clustering programs and found that FlowGrid produces similar clustering results but with substantially less time. For example, FlowGrid is able to complete a clustering task on a data set of 23.6 million cells in less than 12 seconds, while other algorithms take more than 500 seconds or get into error.

FlowGrid is an ultrafast clustering algorithm for large single-cell flow cytometry data. The source code is available at https://github.com/VCCRI/FlowGrid.

## 1. Introduction

Recent technological advancement has made it possible to quantitatively measure the expression of a handful of protein markers in millions of cells in a flow cytometry experiment [11]. The ability to profile such a large number of cells allows us to gain insights into cellular heterogeneity at an unprecedented resolution. Traditionally, cell types are identified based on manual gating of several markers in flow cytometry data. Manual gating relies on visual inspection of a series of two dimensional scatter plots, which makes it difficult to discover structure in high dimensions. It also suffers subjectivity, in terms of the order in which pairs of protein markers are explored, and the inherent uncertainty of manually drawing the cluster boundaries [9]. An emerging solution is to use unsupervised clustering algorithms to automatically identify clusters in potentially multidimensional flow cytometry data.

The Flow Cytometry Critical Assessment of Population Identification Methods (Flow-CAP) challenge has compared the performance of many flow cytometry clustering algorithms [1]. In the challenge, ADIcyt has the highest accuracy but has a long runtime, which makes it impractical for routine usage. Flock [8] maintains a high accuracy and reasonable runtime. After the challenge, several algorithms have been built for flow cytometry data analysis such as FlowPeaks [3], FlowSOM [10] and BayesFlow [6].

FlowPeaks and Flock are largely based on *k*-means clustering. *k*-means clustering requires the number of clusters (*k*) to be defined prior to the analysis. It is hard to determine a suitable *k* in practice. FlowPeaks performs *k*-means clustering with a large initial *k*, and iteratively merges nearby clusters that are not separated by low density regions into one cluster. Flock utilises grids to identify high density regions, which the algorithm then uses to identify initial cluster centres for *k*-means clustering. This grid-based method of identifying high density region allows *k*-means clustering to converge much quicker compared to using random initialisation of cluster centres, and also directly identifies a suitable value for *k*. FlowSOM starts with training Self-Organising Map (SOM), followed by consensus hierarchical clustering of the cells for meta-clustering. In the algorithm, the number of clusters (*k*) is required for meta-clustering.

BayesFlow uses a Bayesian hierarchical model to identify different cell populations in one or many samples. The key benefit of this method is its ability to incorporate prior knowledge, and captures the variability in shapes and locations of populations between the samples [6]. However, BayesFlow tends to be computational expensive as Markov Chain Monte Carlo sampling requires a large number of iterations. Therefore, BayesFlow is often impractical for flow cytometry data sets of realistic size.

These algorithms perform well on the Flow-CAP data sets, but they may not be scalable to larger data sets that we are dealing with nowadays – those with tens of millions of cells. Aiming to quantify cell population heterogeneity in huge data sets, we have to develop an ultrafast and scalable clustering algorithm.

In this paper, we present a new clustering algorithm that combines the benefit of DBSCAN[2] (a widely-based density-based clustering algorithm) and a grid-based approach to achieve scalability. DBSCAN is fast and can detect clusters with complex shapes in the presence of outliers [2]. DBSCAN starts with identifying core points that have a large number of neighbours within a user-defined region. Once the core points are found, nearby core points and closely located non-core points are grouped together to form clusters. This algorithm will identify clusters that are defined as high-density regions that are separated by the low-density regions. However, DBSCAN is memory inefficient if the data set is very large, or has large highly connected components.

To reduce the computational search space and memory requirement, our algorithm extends the idea of DBSCAN by using equal-spaced grids like Flock. We implemented our algorithm in an open source python package called FlowGrid. Using a range of real data sets, we demonstrate that FlowGrid is much faster than other state-of-the-art flow cytometry clustering algorithms, and produce similar clustering results. The detail of the algorithm is presented in the Methods section.

## 2. Methods

The key idea of our algorithm is to replace the calculation of density from individual points to discrete bins as defined by a uniform grid. This way, the clustering step of the algorithm will scale with the number of nonempty bins, which is significantly smaller than the number of points in lower dimensional data sets. Therefore the overall time complexity of our algorithm is dominated by the binning step, which is in the order of *O(N)*. This is significantly better than the time complexity of DBSCAN, which is in the order of *O(NlogN)*. The definition and algorithm are presented in the following subsections.

### 2.1. Definition

The key terms involved in the algorithm are defined in this subsection. A graphical example can be found in Figure 1.

**Figure 1.**
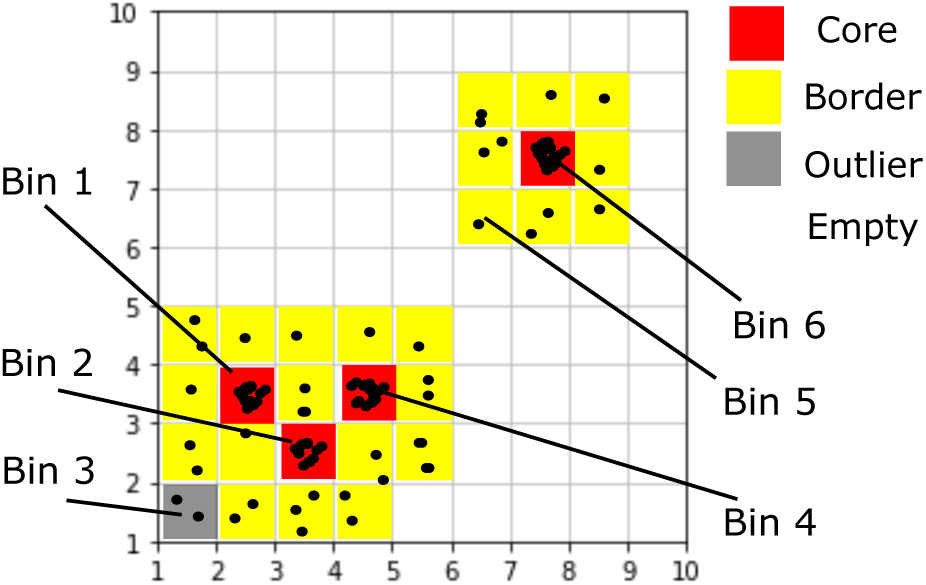
An illustrative example of the FlowGrid clustering algorithm.

- *N*_*bin*_ is the number of equally sized bins in each dimension. In theory, there are (*N*_*bin*_)^*d*^ bins in the data space, where *d* is the number of dimensions. However, in practice, we only consider the non-empty bins. The number of non-empty bins (*N*) is less than (*N*_*bin*_)^*d*^, especially for high dimensional data. Each non-empty bin is assigned an integer index *i* = 1… *N*.
- *Bin*_*i*_ is labelled by a tuple with *d* positive integers *C*_*i*_= (*C*_*i1*_, *C*_*i2*_, *C*_*i3*_, …, *C*_*id*_) where *C*_*i*1_ is the coordinate (the bin index) at dimension 1. For example, if *Bin*_*i*_ has coordinate *C*_*i*_= (2, 3, 5), this bin is located in second bin in dimension 1, third bin in dimension 2 and the fifth bin in dimension 3.
- The distance between *Bin*_*i*_ and *Bin*_*j*_ is defined as

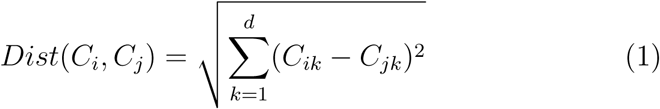
- *Bin*_*i*_ and *Bin*_*j*_ are defined to be directly connected if *Dist*(*C*_*i*_,*C*_*j*_) ⩽ 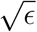, where *ϵ* is a user-specified parameter.
- *Den*_*b*_(*C*_*i*_) is the density of *Bin*_*i*_, which is defined as the number of points in *Bin*_*i*_.
- *Den*_*c*_(*C*_*i*_) is the collective density of *Bin*_*i*_, calculated by

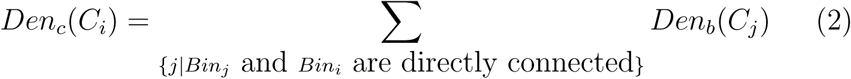
- *Bin*_*i*_ is a core bin if

1. *Den*_*b*_(*C*_*i*_) is larger than *MinDen*_*b*_, a user-specified parameter.
2. *Den*_*b*_(*C*_*i*_) is larger than *ρ*% of its directly connected bins, where *ρ* is a user-specified parameter.
3. *Den*_*c*_(*C*_*i*_) is larger than *MinDen*_*c*_, a user-specified parameter.
- *Bin*_*i*_ is a border bin if it is not a core bin but it is directly connected to a core bin.
- *Bin*_*i*_ is an outlier bin, if it is not a core bin nor a border bin.
- *Bin*_*a*_ and *Bin*_*b*_ are in the same cluster, if they satisfy one of the following conditions:

1. they are directly connected and at least one of them is core bin;
2. they are not directly connected but are connected by a sequence of directly connected core bins.
- Two points are in the same cluster, if they belong to the same bin or their corresponding bins belong to the same cluster.

### 2.2. Algorithm

Algorithm 1 describes the key steps of FlowGrid, starting with normalising the values in each dimension to range between 1 and (*N*_*bin*_ + 1). Then, we use the integer part of the normalised value as the coordinate of its corresponding bin. Then, the SearchCore algorithm is applied to discover the core bins and their directly connected bins. Once the core bins and connections are found, Breadth First Search(BFS) is used to group the connected bins into a cluster. The cells are labelled by the label of their corresponding bins.

## 3. Evaluation

### 3.1. Procedure

FlowGrid aims to be an ultrafast and accurate clustering algorithm for very large flow cytometry data. Therefore, both the accuracy and scalability performance need to be evaluated. The benchmark data sets from Flow-CAP [1], the multi-centre CyTOF data from Li *et al*. [7] and the SeaFlow project [5] are selected to compare the performance of FlowGrid against other state-of-the-art algorithms, FlowSOM, FlowPeaks, and FLOCK. These three In the example in Figure 1, Bin 1, Bin 2, Bin 3 and Bin 6 are core bins as their *Den*_*b*_ are larger than *MinDen*_*b*_ (5 in this example), their *Den*_*c*_ are larger than *MinDen*_*c*_ (20 in this example), and their *Den*_*b*_ are larger than *rho%* (75% in this example) of its directly connected bins. *Dist*(*C*_1_,*C*_2_) = 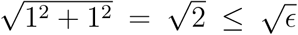 (*ϵ* = 2 in this example), so Bin 1 and Bin 2 are directly connected. 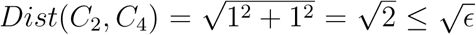, so Bin 2 and Bin 4 are directly connected. Therefore, Bin 1, Bin 2 and Bin 4 are mutually connected, and they are assigned into the same cluster. Bin 5 is not a core bin but is a border bin, as it is directly connected to Bin 6, which is a core bin. Bin 3 is a outlier bin, as it is not a core bin nor a border bin. In practice, *MinDen*_*b*_ is set to be 3, *MinDen*_*c*_ is set to 40 and *rho* is set to be 85.

**input**: *X, N*_*bin*_,*ϵ, ρ, MinDen*_*b*_, *MinDen*_*c*_

**output:** DataLabel

1. Normalise the data *X* ranging from 1 to (*N*_*bin*_ + 1)
2. Assign data into corresponding bins based on the integer of normalised value
3. Identify *S*_*bin*_ as the set of non-empty bins
4. Search the core bins and their directly connected bins by SearchCore
5. Group connected bins into a cluster by Breadth First Search(BFS)
6. Label cells by the label of their corresponding bins

~~~
**Algorithm 1:** FlowGrid
**input**: *S*_*bin*_,*ϵ, ρ, MinDen*_*b*_, *MinDen*_*c*_
**output:** *S*_*core*_, *L*
Initial an empty adjacency list *L*.
*S*_*core*_, *L*
**forall** *Bin*_*i*_ *in S*_*bin*_ **do**
 | **if**  *Den*_*b*_(*i*) >*MinDen*_*b*_ **then**
 |  |  nnBin=radiusNeighbors(*S*_*bin*_, *Bin*_*i*_, *ϵ*)
 |  |  nnCount counting the number of points for each bin in nnBin
 |  |  **if**  *Den*_*b*_(*i*) *is greater than ρ% of nnCount* **then**
 |  |   |  *Den*_*c*_(*i*)= the sum of nnCount
 |  |   |  **if**  *Den*_*c*_(*i*)>*MinDen*_*c*_ **then**
 |  |   |   |  *S*_*core*_ = *S*_*core*_ ∪ {*i*}
 |  |   |   |  mapping *bin*_*i*_ with nnBin in L
 |  |   |   **end**
 |  |   **end**
 | **end
 end**
The input of radiusNeighbors is all non-empty bins, the query bin and the maximum query distance 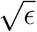. The output is the bins whose distance with the query bin are less than 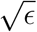 (including the query bin).
~~~

~~~
**Algorithm 2:** SearchCore
algorithms are chosen because they are widely used, are generally considered to be quite fast, and have good accuracy.
Three benchmark data sets from Flow-CAP [1] are selected for evaluation, including the Diffuse Large B-cell Lymphoma (DLBL), Hematopoietic Stem Cell Transplant (HSCT), and Graft versus Host Disease(GvHD) data set. Each data set contains 10-30 samples with 3-4 markers, and each sample includes 2,000-35,000 cells.
The multi-centre CyTOF data set from Li *et al*. [7] provides a labelled data set with 16 samples. Each samples contains 40,000-70,000 cells and 26 markers. Since only 8 out fo 26 markers are determined to be relevant markers in the original paper [7], only these 8 markers are used for clustering.
We also use three data sets from the SeaFlow project [5] and they contain many samples. Instead of analysing the independent samples, we analyse the concatenated data sets as the original paper [5] and these concatenated data
**input**: *S*_*core*_, *S*_*bin*_, adjacency list L
**output**: Bin Label
Label every bin as −1
Index=1
**for** *Bin*_*i*_ *in S*_*core*_ **do**
 | **if** *the laebl of Bin*_*i*_ *is −1* **then**
 |  |  Queue={}
 |  |  Label *Bin*_*i*_ as Index
 |  |  Queue.push(*Bin*_*i*_)
 |  |  **while** *Queue is not empty* **do**
 |  |  |   *Bin*_1_= Queue.pop()
 |  |  |   **forall** *directed connected Bin*_2_ *of Bin*_1_ **do**
 |  |  |    | **if** *the laebl of Bin*_2_ *is −1* **then**
 |  |  |    |  |  Label *Bin*_2_ as Index
 |  |  |    |  | **if** *Bin*_2_ *is core bin* **then**
 |  |  |    |  |  |  Queue.push(*Bin*_2_)
 |  |  |    |  | **end**
 |  |  |    | **end**
 |  |  |   **end**
 |  | **end**
 |  | index=index +1
 | **end
end**
~~~

#### Algorithm 3: Breadth First Search(BFS)

sets contain 12.7, 22.7 and 23.6 millions of cells respectively. Each data sets include 15 features but the original study only uses four features for clustering analysis. The four features are forward scatter (small and perpendicular), phycoerythrin, and chlorophyll (small) [5].

In the evaluation, we treat the manual gating label as the gold standard for measuring the quality of clustering. In the pre-precessing step, we apply the inverse hyperbolic function with the factor of 5 to transform the multi-centre data and the SeaFlow data. As the Flow-CAP and multi-centre CyTOF data contain many samples and we treat each sample as a data set, we run all algorithms on each sample. The performances are measured by the ARI and runtime, which are reported by the arithmetic means (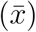) and standard deviation (*sd*). For the Seaflow data sets, we treat each concate nated data set as a data set. In the evaluation, all algorithms are applied on these concatenated data sets.

To evaluate the scalability of each algorithm, we down-sample the largest concatenated data set from the SeaFlow project, generating 10 sub-sampled data sets in which the numbers of cells range from 20 thousand to 20 million.

#### 3.1.1. Performance measure

The efficiency performance is measured by the runtime while the clustering performance is measured by Adjusted Rand Index (ARI). ARI is used to measure the clustering performance. ARI is the corrected-for-chance version of the Rand index [4]. Although it may result in negative values if the index is less than expected, it tends to be more robust than many other measures like F-measure and Rand index.

ARI is calculated as follow. Given a set *S* of *n* elements, and two groups of cluster labels (one group of ground truth label and one group of predicted labels) of these elements, namely *X* = {*X*_1_,*X*_2_, …, *X*_*r*_} and *Y* = *Y*_1_,*Y*_2_,…, *Y*_*s*_}, the overlap between X and Y can be summarized by *n*_*ij*_ where *n*_ij_ denotes the number of objects in common between *X*_*i*_ and *Y*_*j*_: *n*_*ij*_ = |*X*_*i*_ ∩ *Y*_*j*_|.

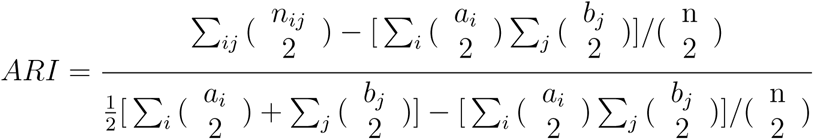

where *a*_*i*_ = Σ_*j*_ *n*_*ij*_ and *b*_*j*_ = Σ_*j*_ *n*_*ij*_

### 3.2. Experimentation

FlowGrid is publicly available as an open source program on github https://github.com/VCCRI/FlowGrid. FlowSOM and FlowPeaks are available as R packages from Bioconductor. The source code of Flock is downloaded from https://sourceforge.net/projects/immportflock/files/FLOCK_flowCAP-I_code. To reproduce all the comparisons presented in this paper, the source code and data can be downloaded from https://github.com/VCCRI/FlowGrid_compare. We run all the experiments on six 2.60GHz cores CPU with 32G RAM.

FlowPeaks and Flock provide automated version without any user-input parameter. FlowSOM requires one user-supplied parameter (*k*, the number of clusters in meta-clustering step). FlowGrid requires two user-supplied parameters (*bin*_*n*_ and *ϵ*). To optimise the result, we try many *k* for FlowSOM and many combinations of *bin*_*n*_ and *ϵ* for our algorithm.

## 4. Results

### 4.1. Performance comparison

Table 1 summaries the performance of our algorithm and three other algorithms - FlowSOM, FlowPeaks, and Flock in terms of runtime. Our algorithm is substantially faster than other clustering algorithms in all the data sets. This improvement of runtime is especially substantial in the Seaflow data sets. FLOCK and FlowPeaks sometimes fail to complete in some of the data sets. In a data set of 23.6 million cells, FlowSOM completes the execution in 572 seconds, whereas FlowGrid completes the execution in only 12 seconds. This is a 50x speed up. Table 2 summaries the clustering accuracy performance. In Flow-CAP and the multi-centre data sets, FlowGrid shares the similar clustering accuracy (in terms of ARI) with other clustering algorithms but in Seaflow data sets, FlowGrid gives higher accuracy than other clustering algorithms.

Figure 2 shows that the clustering results of our algorithm and three other algorithms in a HSCT sample. FlowGrid, FlowSOM and FlowPeaks recover similar number of clusters, and the clustering results are largely similar. Flock generates too many clusters in this case. It is important to note that FlowGrid also identifies cells that do not belong to a main cluster (*i.e*., a high density region). These cells can be viewed as ‘outliers’, and are labelled as ‘−1’ in Figure 2. This is a feature that is not present in other clustering algorithms.

**Figure 2.**
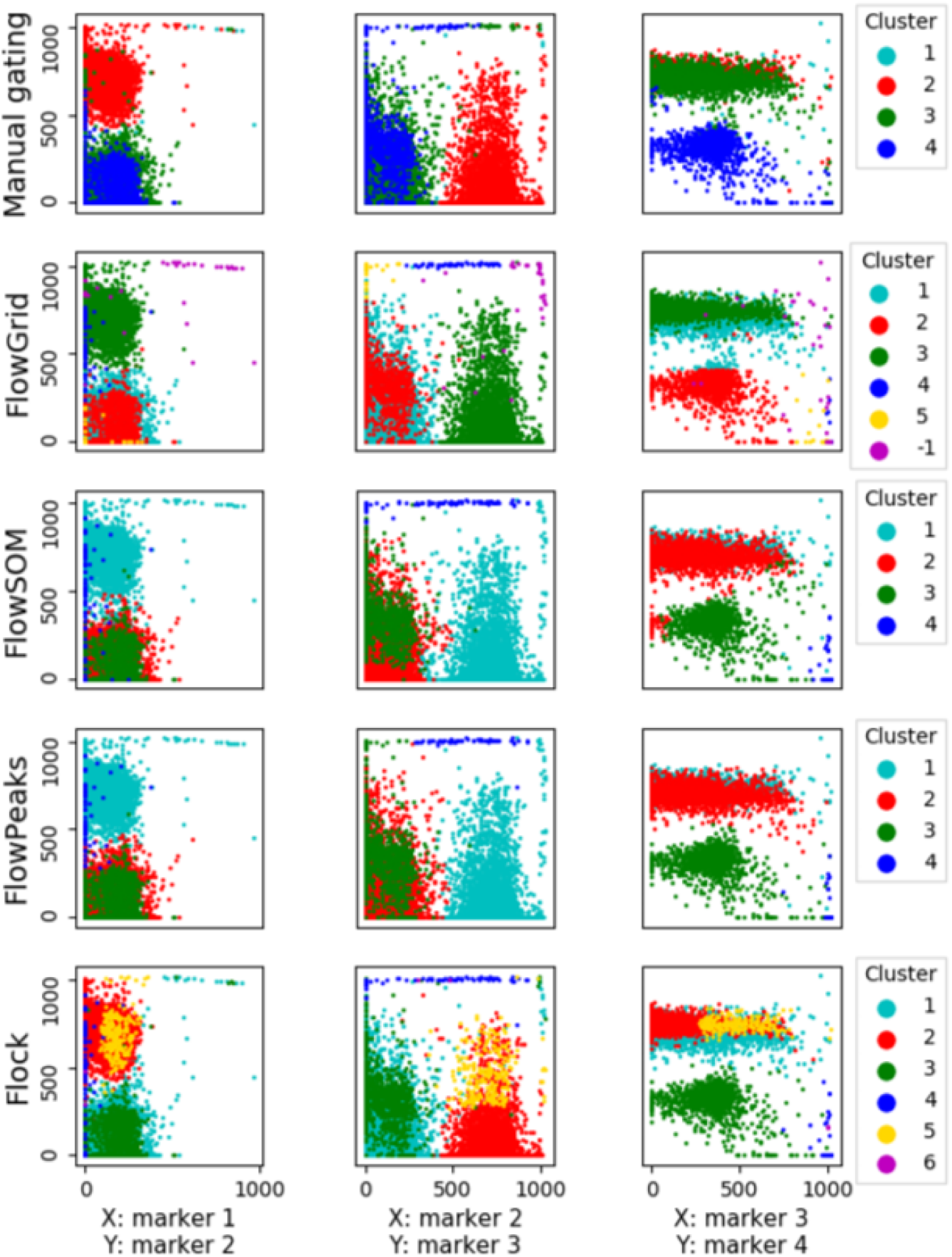
Visual comparison of the clustering performance of FlowGrid, FlowPeaks, Flow-SOM, and Flock using manual gating (top row) as the gold standard.

### 4.2. Scalability analysis

To further evaluate the scalability of the algorithms, we sub-sample one Seaflow data set and the sampled data sets range from 20 thousand to 20 million cells. Figure 3 shows the scalability of our algorithm and three other algorithms. Flock has a low runtime when processing a small data set, but its runtime dramatically increases to 6,640 seconds for a 20 million-cell data set. FlowPeaks and FlowSOM share similar scalability but FlowPeaks is not able to execute 20 million data set. Our algorithm have the best performance in the evaluation as FlowGrid is faster than other algorithm in all the sampled data by an order of magnitude.

**Figure 3:**
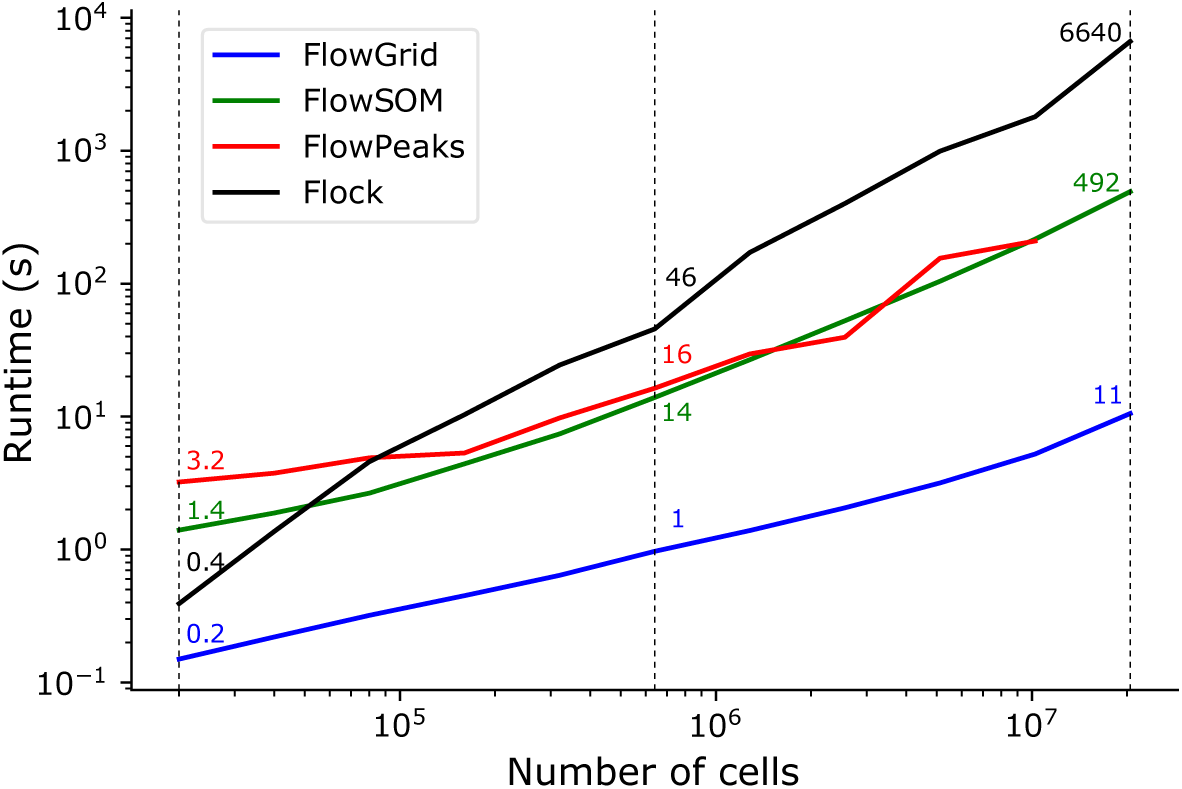
Comparison of the runtime of FlowGrid, FlowPeaks, FlowSOM, and Flock using data sets with different number of cells.

### 4.3 Parameter robustness analysis

Like other density-based clustering algorithm, parameter setting is important. In our experience, *Bin*_*n*_ and *ϵ* are data-set-dependent. We recommend trying out different combinations of *Bin_n_* between 4 and 15, and e between 1 and 5. To pick the best parameter combinations, some prior knowledge is helpful such as the expected number of clusters and the proportion of outliers which should be less than 10% in our experience.

We found that other parameters, namely *MinDen*_*b*_, *MinDen*_*c*_ and *ρ* are mostly robust across a wide range of values.

To demonstrate this robustness, we used the benchmark data sets from Flow-CAP for a parameter sensitivity analysis. For these experiments, we first set 3, 40, 85, 4 and 1 as the default value for *MinDen*_*b*_, *MinDen*_*c*_, *ρ, Bin*_*n*_ and *ϵ*, respectively. In each experiment, we only change one parameter to test its sensitivity to the overall classification result. The performance is measured by ARI and runtime. In the first experiment, we varied *MinDen*_*b*_ from 1 to 50 while fixing other parameters. In the second experiment, we varied *MinDen_c_* ranging from 10 to 300 while fixing other parameters. In the third experiment, we varied *ρ* ranging from 70 to 95 while fixing other parameters.

Figure 4 demonstrates that the clustering accuracy and runtime are largely insensitive to *MinDen*_*b*_, *MinDen*_*c*_ and *ρ* across a large range of parameter values. The experiments are applied to all the benchmark data sets from Flow-CAP and similar results are observed in all the benchmark data sets. In our experiments, when *MinDen*_*b*_, *MinDen*_*c*_ and *ρ* are set to be 3, 40 and 85 respectively, FlowGrid maintains good clustering performance and excellent runtime. They are therefore set as the default parameters for FlowGrid.

**Figure 4.**
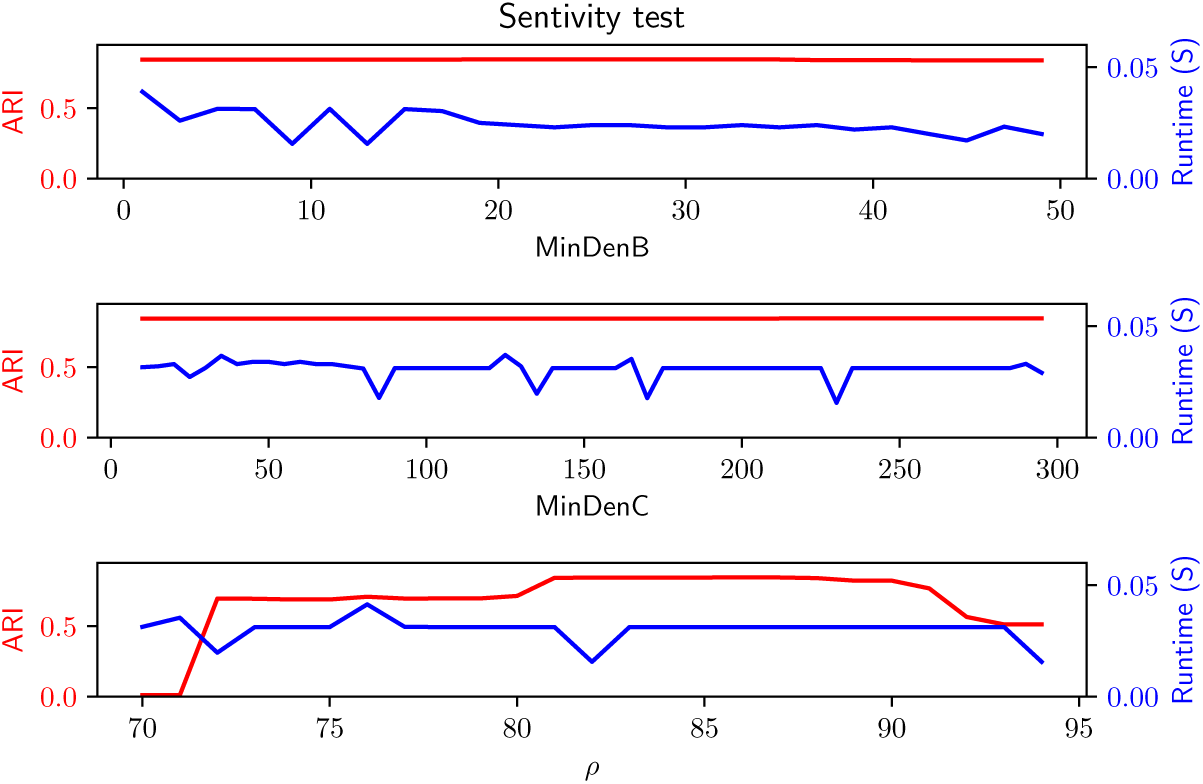
Sensitivity analysis of three different parameters on clustering accuracy (as measured by adjusted rand index; ARI) and runtime (seconds).

## 5. Discussion

In this paper, we have developed an ultrafast clustering algorithm, FlowGrid, for single-cell flow cytometry analysis, and compared it against other state-of-the-art algorithms such as Flock, FlowSOM and FlowPeaks. FlowGrid borrows ideas from DBSCAN for detection of high density regions and outliers. It does not only perform well in the presence of outliers, but also have great scalability without getting into memory issues. It is both time efficient and memory efficient. FlowGrid shares similar clustering accuracy with state-of-the-art flow cytometry clustering algorithms, but it is substantially faster than them. With any given number of markers, the runtime of FlowGrid scales linearly with the number of cells, which is a useful property for extremely large data sets.

*MinDen*_*b*_ and *MinDen*_*c*_ are density threshold parameters to reduce the search area of high density bin. If the parameters are set very low, the run-time may fractionally increase but the accuracy is not likely to be affected. However, if the parameters are set very high, the runtime will also fractionally decreases but it may lead to the separation of real clusters and create spurious outliers. In any case, we showed that the performance of FlowGrid is generally robust against changes in *MinDen*_*b*_, *MinDen*_*c*_ and *ρ*.

The current implementation of FlowGrid is already very fast for most practical purposes. In the future, if the data size grows even larger, it is possible to further speed up FlowGrid by parallelising the binning step of the algorithm, which is currently the most computationally intensive step of the algorithm.

